# Global inhibition in head-direction neural circuits: a systematic comparison between connectome-based spiking neural circuit models

**DOI:** 10.1101/2023.02.03.526928

**Authors:** Ning Chang, Hsuan-Pei Huang, Chung-Chuan Lo

**Author notes:** equal contribution (Ning Chang, Hsuan-Pei Huang).

## Abstract

The recent discovery of the head-direction (HD) system in fruit flies has provided unprecedented insights into the neural mechanisms of spatial orientation. Despite the progress, the neural substance of global inhibition, an essential component of the HD circuits, remains controversial. Some studies suggested that the ring neurons provide global inhibition, while others suggested the Δ7 neurons. In the present study, we provide evaluations from the theoretical perspective by performing systematic analyses on the computational models based on the ring-neuron (R models) and Δ7-neurons (Delta models) hypotheses with modifications according to the latest connectomic data. We conducted four tests: robustness, persistency, speed, and dynamical characteristics. We discovered that the two models led to a comparable performance in general, but each excelled in different tests. The R Models were more robust, while the Delta models were better in the persistency test. We also tested a hybrid model that combines both inhibitory mechanisms. While the performances of the R and Delta models in each test are highly parameter-dependent, the Hybrid model performed well in all tests with the same set of parameters. Our results suggest the possibility of combined inhibitory mechanisms in the HD circuits of fruit flies.

## Introduction

The head-direction (HD) system is a set of neural circuits that encode the head direction of an animal with respect to a reference point, e.g., a landmark. The HD system provides vital information for animal navigation and has been extensively studied in rodents (Taube et al. 1990; Muller et al. 1996; Taube 2007). To elucidate the computational principles of the HD system, several studies have proposed high-level computational models (Skaggs et al. 1995; Redish et al. 1996; Zhang 1996; Goodridge and Touretzky 2000; Xie et al. 2002). However, the detailed neural circuit structure and the underlying mechanisms of the HD system only became available through recent studies in the central complex of fruit flies, *Drosophila melanogaster*. The central complex is a structure in the center of arthropod brains and is known to perform functions related to locomotion, high-level behavioral control, and sensory integration (Strauss and Heisenberg 1993; Strauss 2002; Heinze and Homberg 2007; Ueno et al. 2012; Weir and Dickinson 2015). A recent study discovered that neurons in the ellipsoid body, a neuropil in the central complex, encode the head direction (Seelig and Jayaraman 2015). Furthermore, analyses of the connectome in this region provided strong evidence of the existence of attractor circuits, a common hypothesis of many earlier computational models of the rodent HD system (Lin et al. 2013; Wolff et al. 2015; Chang et al. 2017; Turner-Evans et al. 2020; Hulse et al. 2021).

Based on the recent empirical studies on the central complex, several computational models were proposed (Givon et al. 2017; Kakaria and de Bivort 2017; Cope et al. 2017; Turner-Evans et al. 2017; Su et al. 2017; Stone et al. 2017; Pisokas et al. 2020; Han et al. 2021; Goulard et al. 2021; Lazar et al. 2021), and some of them focused on demonstrating how the head direction is encoded through the attractor dynamics (Kakaria and de Bivort 2017; Su et al. 2017; Pisokas et al. 2020; Han et al. 2021). The basic attractor dynamics require two major components: excitatory neurons that form local recurrent excitation and inhibitory neurons that provide feedback global inhibition to the excitatory neurons (Fig. 1a, left). Although most models suggest that the EPG (or E-PG) neurons, which project from the ellipsoid body to the protocerebral bridge and the gall, take part in providing local recurrent excitation, the assumption of the neurons that provide the global inhibition differs. There are two major hypotheses: the ring neurons (Su et al. 2017; Han et al. 2021) and the Δ7 neurons (Kakaria and de Bivort 2017; Pisokas et al. 2020). The ring neurons are a group of GABAergic projection neurons with many subtypes. They are recognized by the ring-shaped axonal domains innervating all wedges in the ellipsoid body and providing a global inhibition on the HD circuits. In contrast, the dendritic domains are located in different neuropils, mostly in BU (bulb, or lateral triangle) but also in other regions such as LAL (lateral accessory lobe) or CRE (crepine) (Young and Armstrong 2010; Hulse et al. 2021). Δ7 neurons are a set of intrinsic neurons of PB (protocerebral bridge). They are likely to be glutamatergic (Turner-Evans et al. 2020) but are shown to functionally inhibit EPG neurons (Franconville et al. 2018). Each Δ7 neuron projects its dendritic and axonal domains to several PB glomeruli, providing a patterned (or patched) inhibition. But all Δ7 neurons form a tiling pattern that covers the entire PB and also provides a global inhibition on the HD circuits.

**Fig. 1.**
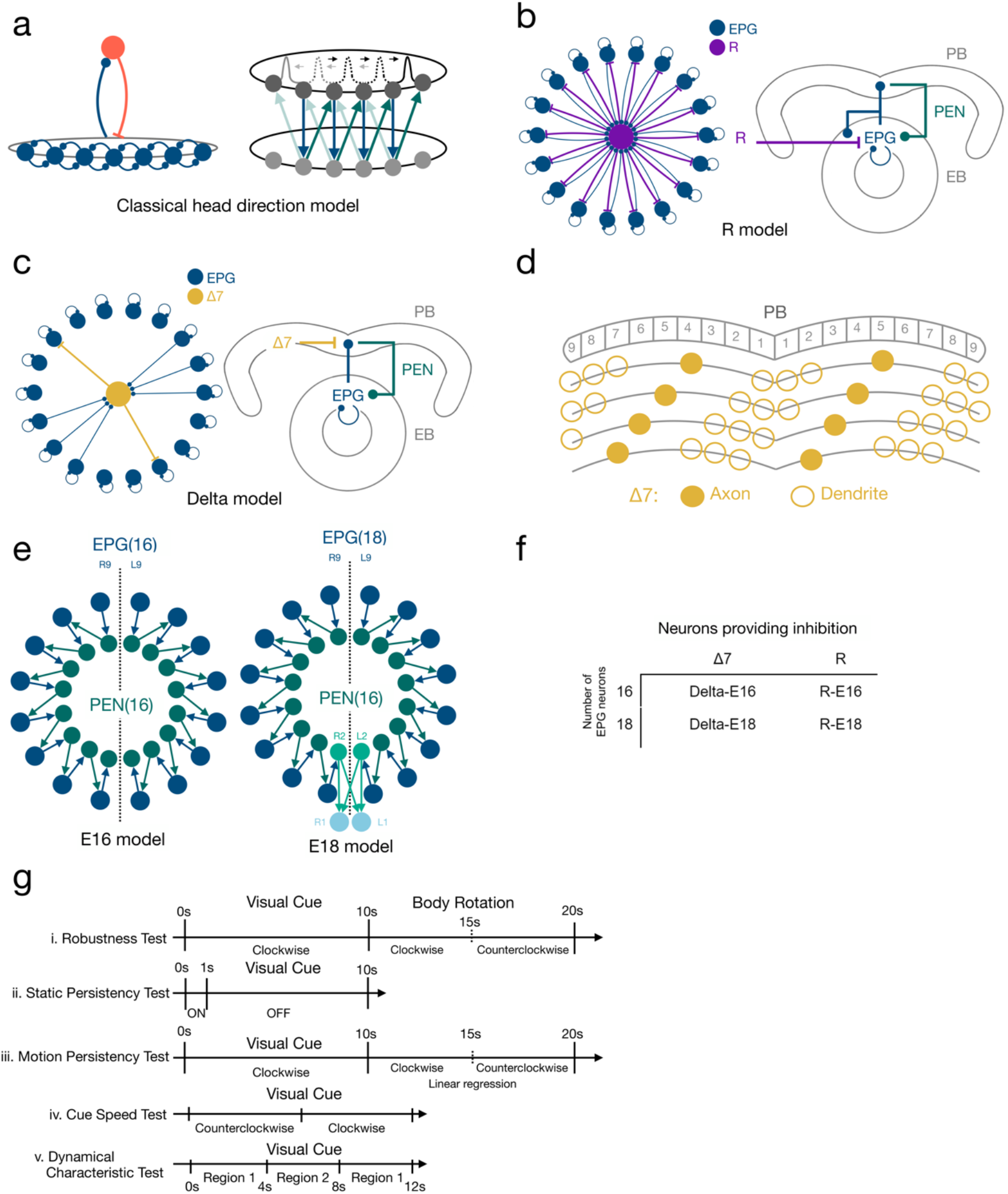
The circuit models and the test protocols. **a** Schematics of the classical head-direction (HD) neural circuits. Left: The circuit diagram of the core component of HD circuits, the ring attractor network. The network consists of excitatory neurons (dark blue), which form locally recurrent excitation, and inhibitory neurons (red), which provide global feedback inhibition. Right: The HD circuits consist of two layers. The top layer is a ring-attractor network that encodes the head direction. The bottom layer receives input from the top layer and feeds back to the neighboring neurons in the top layer, forming shifter circuits. **b** The R class models. The ring (R) neurons provide global inhibition to the attractor network. Left: the attractor circuits of the models. Right: the innervation sites of each neuron type. EB: ellipsoid body. PB: protocerebral bridge. **c** Same as in B but for the Delta class models, in which the Δ7 neurons provide global inhibition. In both classes of the models, EPG neurons form local excitation, and PEN neurons constitute shifter circuits. **d** Four example Δ7 neurons showing the innervation pattern of the inhibitory neurons in the Delta version. Each Δ7 neuron only innervates a subset of glomeruli in PB. However, all eight Δ7 neurons collectively provide global inhibition that covers the entire PB. **e** We also investigate another two variants of the model: The E16 (left) and E18 (right), containing 16 and 18 EPG neurons, respectively. The two extra EPG neurons in the E18 version are indicated in light blue. **f** The four variants of the HD circuits investigated in the present study and their naming. **g** The protocols for testing (from top to bottom) robustness, persistency (static and motion), speed, and dynamical characteristics.

The corresponding modeling studies showed both types of inhibitory neurons to work in the HD system (Kakaria and de Bivort 2017; Su et al. 2017; Pisokas et al. 2020; Han et al. 2021). The distinct innervation patterns of these two types of inhibitory neurons pose two critical questions: Do they affect the dynamics of the HD circuits differently? If they do, which one provides a better mechanism of global inhibition?

In the present study, we systematically investigated these questions. However, instead of using the previously proposed models straight from their original studies (Kakaria and de Bivort 2017; Su et al. 2017), we modified the models based on the recently released and electronic-microscopy-based connectomic data (Scheffer et al. 2020; Turner-Evans et al. 2020; Hulse et al. 2021) while preserving the basic hypothesis of the global inhibition in each model. The latest connectomic data indicate different connection patterns in several neuron types in the HD circuits from the original modeling papers (see Methods and Materials for detail). We incorporate these updated connections, which led to four models, or four variants of the HD circuits. We tested the four models by sweeping through a wide range of parameter space and investigated four properties: robustness, persistency, speed, and dynamical characteristics.

## Methods and Materials

### Model Network Construction

#### Models of neuron and synapse

In the proposed neuron network models, simulation for each neuron is based on the leaky integrate-and-fire (LIF) model with conductance-based synapses described in the previous study (Su et al. 2017; Huang et al. 2019). We chose a spiking neuron model with biophysically realistic synapse models over the simpler firing-rate-based models because a spike-based model reveals several challenges posed by real nervous systems and can be used to explore more detailed neural mechanisms than the rate-based models do. For example, dynamical instability caused by sparse spike inputs cannot be investigated by rate-based models because their synaptic inputs are always continuous. Hence the spiking neuron model is a reasonable choice.

In the present models, the LIF model is given by

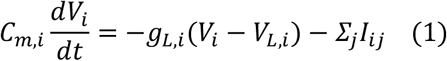

where *C_m_* is the membrane capacitance (=0.1nF for all neurons), *g_L_* = *C_m_*/*τ_m_* is the membrane conductance and is set to make the membrane time constant *τ_m_* equal to 15 ms for each neuron, *V_L_* (= −70mV) is the resting potential, *I_ij_* is the synaptic current elicited by spike input from neuron *j*. The synapses are conductance-based, and three types are modeled in the present study: glutamatergic (via NMDA receptor), cholinergic (Ach), and GABAergic (GABA_A_). The synaptic currents induced by cholinergic, AMPA, and GABAergic receptors are given by

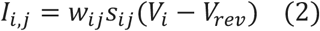

and as for NMDA receptors, the synaptic current is

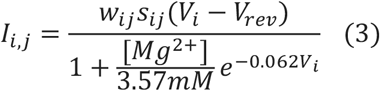

where *w_ij_* is the synaptic conductance (equivalent to synaptic weight), *s_ij_* is the gating variable, *V_rev_* is the reversal potential, which is set to 0mV for all excitatory synapses (including NMDA, and Ach) and is set to −70mV for the inhibitory GABA_A_ synapses. [Mg^2+^] is the extracellular magnesium concentration. The gating variable *s_ij_* describes the activation of channels due to spike input and is given by

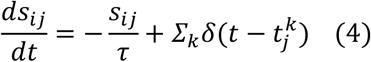

for GABA_A_ and Ach receptors, and by

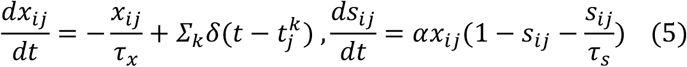

for NMDA receptors, where *τ* is the time constant (5ms, 20ms, and 100ms for GABA_A_, Ach, and NMDA receptors), *δ* is the delta function, 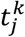 is the time of the *k-th* presynaptic spike from neuron j, *α*(=0.6332) is a factor for adjusting the increment of the NMDA gating variable, or receptor activation rate.

All the neuron and synaptic receptor-related parameters described in this section are fixed in the present models. The value of the membrane time constant (*τ_m_*=15ms) is determined based on the electrophysiological study of the DM1 projection neurons of the fruit fly antennal lobe (Gouwens and Wilson 2009). Other parameters are adopted from the typical values used by other studies or by us in the neural network models of fruit flies or other species (Wang 2002; Lo and Wang 2006; Huang et al. 2019).

#### The network models

In the present study, we constructed two classes of the HD circuit models with one (R class) based on Su et al. 2017 and Han et al. 2021 and another (Delta class) based on Kakaria and de Bivort 2017 and Pisokas et al. 2020. Each class of models included three types of neurons, EPG, PEN, and R (for the R class only) or Δ7 (for the Delta class only). EPG neurons, or the compass neurons, support attractor dynamics and form localized activity, or an activity bump, that encodes the head direction (Clandinin and Giocomo 2015; Seelig and Jayaraman 2015; Turner-Evans and Jayaraman 2016; Kim et al. 2017). PEN neurons form the shifter circuits, and they can move the activity bump clockwise or counterclockwise when the body rotates in the absence of visual input, i.e., in darkness (Fig. 1a, right). The global inhibition, which is essential for the formation of activity bump, is provided by the GABAergic ring (R) neurons (in the R class models) or by the Δ7 neurons (in the Delta class models). Each neuron class (EPG, PEN, R, and Δ7) exhibits a specific innervation pattern at the neuropil level and can be further divided into several types (EPG-L1, EPG-L2, for example), which innervate different subregions of the neuropils. Based on the connectomic data, 1-5 neurons were identified for each type (Scheffer et al. 2020). Therefore, we included three identical neurons for each type in the present models.

Instead of using the exact circuit structures proposed in the original studies, we revised the circuits of each model based on the recently released connectome of the central complex (Scheffer et al. 2020; Hulse et al. 2021). Several changes were made: (1) In the original R class and Delta class models, the local recurrent excitation was hypothesized to be carried out by the EPG← →PEG loops. PEG is a set of excitatory neurons projecting from PB to EB and the Gall. However, the connectomic data indicate very weak PEG→EPG connections. Instead, there are strong self-recurrent connections within the EPG neuronal group. Therefore, we removed EPG← →PEG loops and replaced them with EPG← →EPG loops in both classes of models (Fig. 1b & 1c). (2) In the original Delta class models, each Δ7 neuron innervates all 18 glomeruli in the PB with two axonal and 16 dendritic connections. However, the connectomic data (Scheffer et al. 2020) do not support a broad dendritic innervation in PB. With a careful analysis (Supplemental Fig. 1), we determined that dendrites of each Δ7 neuron only innervate 6-8 PB glomeruli (Fig. 1d). (3) A recent study (Hulse et al. 2021) revealed a total of 18 EPG neuron types, EPG-L1 to L9 and EPG-R1 to R9. Except for EPG-L1 and R2, each of the remaining 16 EPG neurons provides inputs to one of the 16 PEN neurons and receives PEN inputs from neighboring PEN neurons, forming perfect clockwise or counterclockwise shifter circuits (Fig. 1e, left). In contrast, the EPG-L1 and R1 neurons break this pattern by not projecting to any PEN neuron (Fig. 1e, right). Because it is not clear whether these two “atypical” neurons participate in the head-direction function, we created two variants of the models, one with EPG-L1 and R1 neurons (named E18) and one without (E16) for each of the R and Delta model classes. Therefore, we investigated four basic models in the present study: R-E18, R-E16, Delta-E18, and Delta-E16 (Fig. 1f). The connection tables and innervation tables of the four models are shown in supplemental Fig. 2 and 3, respectively. We also investigated the Hybrid model, which combined R-E16 and Delta-E16.

Previous studies reported that the PEN class neuron consists of two subtypes: PEN_a and PEN_b (also named PEN1 and PEN2, respectively)(Green et al. 2017; Hulse et al. 2020). The functional differences between both subtypes are unclear. In the present study, we regarded the PEN neurons as the PEN_a subtype because PEN_a dendrites innervate EPG neurons directly in PB, while PEN_b neurons only contact EPG neurons indirectly via PEG neurons (Turner-Evans et al. 2020), which the present models do not need.

As mentioned earlier, each Δ7 neuron only innervates a subset of the PB glomeruli. However, the whole population of Δ7 neurons tiles the PB and forms a complete and homogenous coverage, except for three atypical Δ7 neurons, L4R6_R, L6R4_L, and L7R3_L. These three neurons do not follow the order of the innervation pattern exhibited by other Δ7 neurons (Supplemental Fig. 4). We have tested the Delta models by including the three atypical Δ7 neurons, but the models failed to work completely. Therefore, in the present study, we excluded L4R6_R, L6R4_L, and L7R3_L from the two Delta models (Delta-E18 and Delta-E16).

Note that although the network structure of each model was determined by the connectomic data, we kept the synaptic weight (*w_ij_* in Equations 2 & 3) of each connection as a tuning parameter. The weight of each synapse in the models is a product of two variables: factor and base. The base value (*K*) defines the basic weight of synapses between each pair of neuron classes (PEN to EPG, for example) and is a tunable parameter in the present study (see Fig. 2c and supplemental Fig. 5 for tuning ranges). The factor is used to scale the weight values within each pair of neuron classes and is not tunable (Supplemental Fig. 2). For example, the factor is 2 for PEN-L3→EPG-R8 but is 1 for PEN-L3→PEG-R7, and all PEN to EPG connections share the same base value, which ranged from 2 to 25 when we swept through the parameter space and looked for usable parameters. The connectomic data provide information about the number of synapses formed between each pair of neurons (Scheffer et al. 2020; Hulse et al. 2020). Although it is tempting to use such information to indicate the synaptic weights, the synaptic numbers revealed in the connectomic data are highly variable, even within the same types of connections. For example, the mean synaptic numbers between EPG and PEN_a neurons vary from 1 (averaged over three EPG_L7 neurons → one PEN_a_L2 neuron) to 45 (averaged over two EPG_L1 neurons → one PEN_a_L2 neuron). All tested models failed if we simply set the synaptic weights proportional to these numbers.

**Fig. 2.**
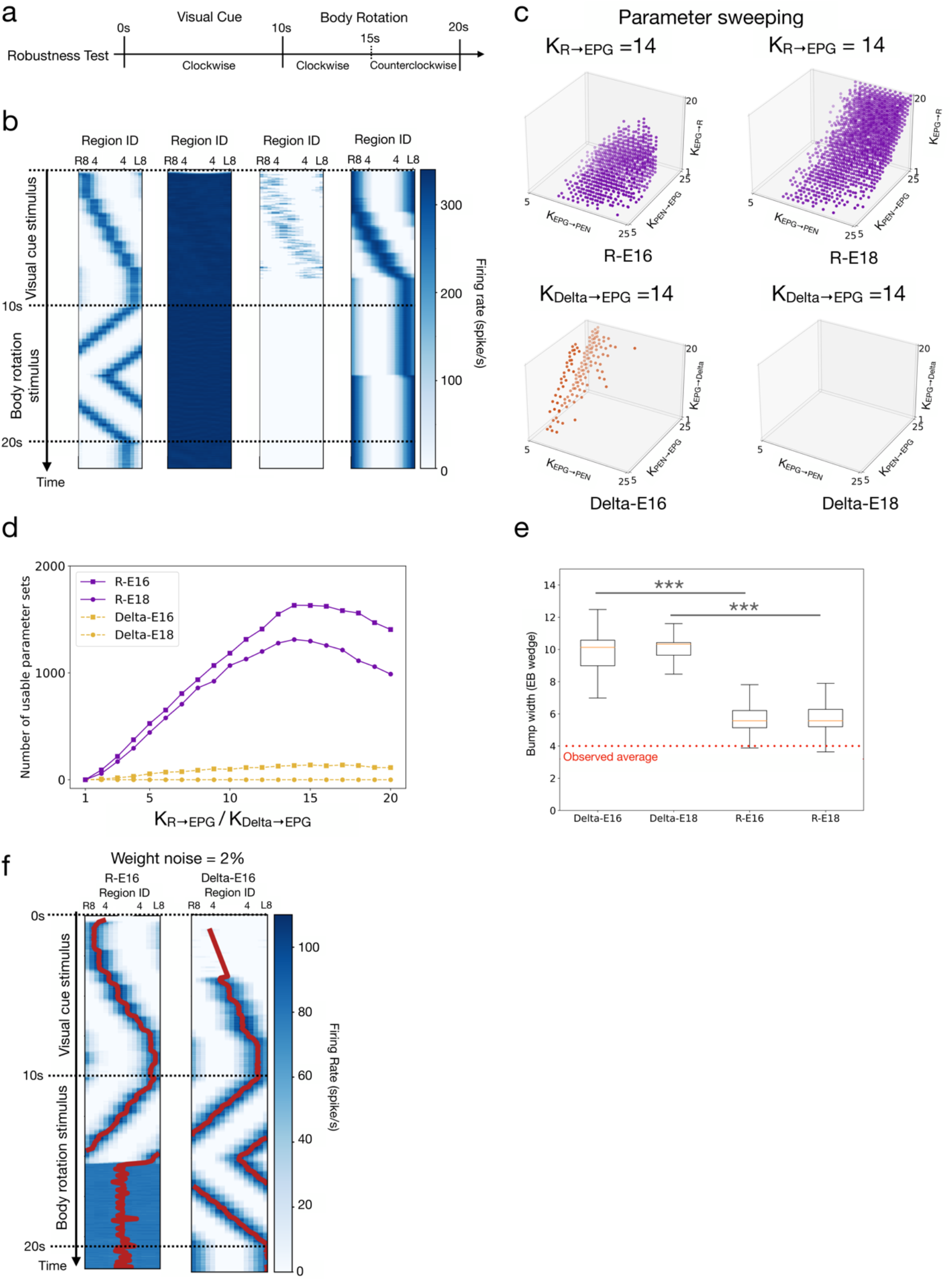
Robustness of the models. **a** The protocol for the robustness test (also shown in Fig. 1g**). b** Examples of one successful trial (left, R-E16) and three failed trials (right three, front left to right: R-E16, R-E16, Delta-E18). The successful trial produced a stable activity bump that tracked visual cue movement and body rotation, while the failed trials did not. **c** We swept through four-dimensional parameter space and marked the parameter sets that led to successful trials (orange and purple dots). The panel shows, for each model, a three-dimensional slice in the four-dimensional sweeping: (top left) R-E16, K_R→EPG_=14, (top right) R-E18, K_R→EPG_=14, (bottom left) Delta-E16, K_Delta→EPG_=14, and (bottom right) Delta-E18, K_Delta→EPG_=14. K is the weight base, which needs to be multiplied by the factor shown in supplemental Fig. 2 to become the synaptic weight. **d** Robustness (defined by the number of usable parameter sets) as a function of the Δ7/R→EPG synaptic strength for all models. The R models were more robust than the Delta models **e** The widths (FWHM) of the bumps in all four models. The red dashed line shows the experimental bump width data from (Kim et al., 2017). Due to the innervation patterns of the Δ7 neurons (Fig. 1d), the mean bump widths of the Delta models were significantly larger than the R models (t-test, ***=p<0.001). An EB wedge is 22.5° (or 0.125π) wide. **f** Example trials of the R-E16 and Delta-E16 models with heterogeneity in the synaptic weights of the EPG→R/Delta and Delta/R→EPG connections.

### Simulation Protocols

#### Estimation of calcium activity and the activity bump

In the present study, the outputs of the models are spike trains, which are different from the calcium activities observed in most experimental studies. To compare the modeled and observed EPG activity in each EB wedge, we followed the method used by Su et al. 2017. Specifically, for each EB wedge *w*, we calculated the population mean activity *r_w_*(*t*) of the EPG neurons by

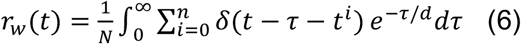

where *N* is the number of EPG neurons projected to the wedge *w*, n is the number of total spikes generated by these *N* neurons, *t^i^* is the time of the *i-th* spike, *δ* is the delta function, and *d* (=721.5ms) is the exponential kernel indicating a 500ms half-life, mimicking the hundred seconds of the half-life of the calcium indicator GCaMP6f (Chen et al. 2013). We further estimated the activity bump by performing Gaussian fitting to *r_w_*(*t*) over *w* for each timestep *t*. The resulting Gaussian function was treated as the representation of the activity bump. The peak position, height, and width (FWHM, Full width at half maximum) of the Gaussian function were used in the subsequent analyses.

#### Visual cue and body rotation

As demonstrated in previous studies (Seelig and Jayaraman 2015; Green et al. 2017), the activities of EPG neurons in EB and PEN neurons in PB encode the azimuthal position of visual stimuli. For simplicity, we mapped the 16 EB wedges to the 360° horizontal visual space as described in the previous study (Su et al. 2017). We activated the corresponding EB wedge to simulate the presentation of the visual cue. The present study used two types of stimulations, visual cue and body rotation, depending on the purpose of the tests. For the visual cue stimulation, we activated the corresponding EB wedges by stimulating the PEN neurons, which innervate the EB wedges. The activation is done with a spike rate of 50 Hz and maximum conductance of 2.1ns on the cholinergic receptors. Note that the same effect can be achieved by stimulating the EPG neurons. However, both methods are equivalent from the modeling perspective due to the direct connections from PEN to EPG neurons. The second type of stimulation represents body rotation and is used to move the activity bump in the darkness without any visual input. This is done by activating unilateral PB via spike inputs (2,210Hz, with a conductance of 0.3 ns on the NMDA receptors) to PEN L2-L9 or R2-R9 neurons, which constitute the shifter circuits that move the activity bump clockwise or counterclockwise, respectively.

#### Robustness Test

In the robustness test, we swept through a wide range of synaptic weight bases (*K*) to determine the working parameter space of each model. The test included four types of synapses, *K*_EPG→PEN_, *K*_PEN→EPG_, *K*_Delta→EPG_ (or K_R→EPG_), *K*_EPG→Delta_ (or *K*_EPG→R_), and the bases ranges from 5ns to 25ns with a step of 1 for the first two types of synapses and 1ns to 20ns with a step of 1 for the last two. Therefore, a total of 176,400 sets of weights were tested for each model. We ran one trial for each set of weights, and each trial consisted of a 10s period of visual cue stimulation followed by a 10s period of body rotation stimulation. The cue moved 45 degrees per second in the visual cue period, while no visual cue was presented in the body rotation period. The 10s body rotation period was further separated into a 5s counterclockwise rotation (stimulating PEN R2-R9) followed by a 5s clockwise rotation (stimulating PEN L2-L9) (Fig. 1gi). For a given parameter set, a trial was considered as failed if it met any of the following four conditions. First, the amplitude of the activity bump was smaller than 1 spike/s for more than 10 ms (diminished bump condition). Second, the width of the activity bump was larger than 360° for over 10 ms (spread bump condition). Third, the bump failed to move in the body rotation period (11-20s) (immovable bump condition). Fourth, the Gaussian fitting failed to return any result for more than 5 ms (no bump condition).

#### Persistency Test

The persistency test is separated into two sub-tests: the static and motion persistency tests.

The static persistency test examined how well the activity bump maintains a static location after the visual cue stimulus is turned off. The test started with a 1s static visual cue stimulation followed by a 9s total darkness (visual cue off) without any body rotation stimulation (Fig. 1gii). A trial was considered as successful if no “diminished bump” and “spread bump”conditions occurred (see **Robustness Test** above for the definition of the two conditions). We performed 1000 trials for each model with randomly selected usable parameter sets for each trial. For successful trials, we further analyzed how steady the bump was in the 9s dark period by computing the standard deviation of the bump peak from the original visual cue location.

The motion persistency test used the same test protocol as the robustness test (Fig. 1giii). In the body rotation period, we expect that an HD model’s bump first shifts linearly in the counterclockwise direction for 5s and then reverses the direction for another 5s. To evaluate how persistent the bump exhibits this linear movement, we performed linear regression for the trajectory of the bump peak separately for the first 5s and the last 5s of the body rotation period. We used the mean *R*^2^ values of the two regressions as the measure of persistency. The mean *R*^2^ is between 0 and 1, with 0 representing no detectable bump movement while 1 representing perfect linear movement.

#### Speed Test

The speed test consisted of only a visual cue period. In each trial, the visual cue moved for one round (360°) in the counterclockwise direction, followed by the clockwise direction for another run. (Fig. 1giv). To evaluate how fast the activity bump can follow the moving cue, we tested the following cue speeds: 0.25, 0.28, 0.312, 0.35, 0.42, 0.5, 0.625, 0.83, 1.25, and 2.5 *πrad/s*. For the last two speeds, we allowed additional 3 and 7 rounds in each direction, respectively, so that the trials lasted long enough for the analysis. We randomly selected seven usable parameter sets for each model and simulated 1000 trials for each parameter set in each speed condition. We followed the criteria stated in **Robustness Test** except that the “diminished bump condition” is changed from 10ms to 5ms. We classified each trial as successful or failed and then calculated the success rate by dividing the number of successful trials by the total number of trials. Finally, we computed the mean success rate over the seven parameter sets for each speed condition and each model. The seven parameters are manually picked to be uniformly distributed in the parameter space to ensure that the results reflect the overall performance from an unbiased sampling of the useable parameters.

#### Dynamical Characteristics Test

In the dynamical characteristics test, we tested whether the activity bump “jumps” to the new location accordingly when the visual cue suddenly shifts from the original location to the new location. The bump jumping usually occurs when the two cue locations are far from each other (Kim et al. 2017). To this end, we selected two EB regions that were 180° apart, and the visual cue switched between the two regions twice in each trial (Fig. 1gv). To identify the trials that exhibited successful bump jumping, we checked whether a discontinuity of the active bump occurred immediately following each cue switch event (5s~5.5s and 8s~8.5s) by checking if “no bump condition” was detected during these two switching periods. The success rate is calculated by dividing the number of successful trials by the total number of trials.

## Results

### The models

In the present study, we systematically tested four models, or four variants of the HD circuits: R-E16, R-E18, Delta-E16, and Delta-E18 (see Materials and Methods). All four models feature the basic attractor network structure and the shifter circuits (Fig. 1a). The differences lie in the global inhibition mechanism (R or Delta) and the number of EPG neurons (E16 or E18). The R models use the ring neurons as the neural substrate for the global inhibition, while the Delta models use the Δ7 neurons. These two types of inhibitory neurons innervate the HD circuits differently (Fig. 1b-d). The E16 and E18 models include 16 or 18 EPG neurons, respectively. The EPG neurons in the E16 models form one-to-one feedback excitation with the 16 PEN neurons, while the two extra EPG neurons in the E18 models receive input from PEN but do not provide feedback (Fig. 1e, Supplemental Fig. 2). We tested all four models for their robustness, persistency, speed, and dynamical characteristics (Fig. 1g).

### Robustness

A model is considered robust if it is not sensitive to the selection of parameters. We tested whether an HD model produced an activity bump and tracked the head direction with a given parameter set. A parameter set that did not lead to a “failed trial” (as defined in Materials and Methods) is referred to as a “usable parameter set” (Fig. 2b). Next, we repeated the test by sweeping through a four-dimensional parameter space, spanned by the four types of synaptic weight bases: K_Delta→EPG_ (or K_R→EPG_), K_EPG→PEN_, K_PEN→EPG_, K_EPG→Delta_ (or K_EPG→R_). We visualized the result by marking the usable parameter sets in each three-dimensional slice, given by a specific K_Delta→EPG_ (or K_R→EPG_) value, in the four-dimensional parameter space (Fig. 2c; Supplemental Fig. 5). A model is more robust than others if it has more usable parameter sets. By counting the number of usable parameter sets in each slice, we discovered that R-E16 and R-E18 are much more robust than Delta-E16, and we cannot find any usable parameter for Delta-E18 (Fig. 2d). We noted that the Delta-E18 model failed mainly due to the inability to maintain a moving bump during the body rotation period in the absence of the visual cue. However, Delta-E18 still worked when a visual cue was presented. Therefore, we still included Delta-E18 in the tests which involved visual stimulation. Note that due to the difference in the inhibition mechanism, the mean bump width of the Delta models was significantly larger than that of the R models (1.23π for Delta-E16, 1.26π for Delta-E18, 0.73π for R-E16 and 0.74π for R-E18). The experimentally observed value was ~0.5π (Seelig and Jayaraman 2015; Kim et al. 2017) (Fig. 2e). Finally, we tested how robust the R-E16 and Delta-E16 models are against heterogeneous weight distribution. We added a Gaussian distributed noise to each weight in the EPG→R/Delta and R/Delta→EPG connections and counted the success rates in 100 trials of the Robustness test. As expected for the continuous attractor networks, both R-E16 and Delta-E16 models do not tolerate heterogeneous synaptic weights. Both models failed to work in most trials with a noise of merely 2% of the weights (success rate: Delta-E16 = 0.34, R-E16 = 0.37) (Fig. 2f).

### Persistency

Next, we tested how accurately the activity bump tracks the head direction. When a visual landmark was presented, we observed that the bump could always accurately follow the visual cue, whether persistent or shifting. Therefore, here we focused on the darkness condition, where the bump should remain persistent and only move when the shifter circuits are activated. We first tested how steady the bump is in the absence of light with a fixed head direction (Fig. 3b). We counted the percentage of successful trials (see Materials and Methods) and discovered that the two Delta models performed slightly better than the two R models by exhibiting ~8% more successful trials (Fig. 3c). We further compared the drift of the bump from the cue position by computing the standard deviation between the bump peak and the original cue position. Again, the Delta models performed significantly better than the R models by exhibiting smaller deviations (7.65° for Delta-E16, 2.70° for Delta-E18, and 22.88° for both R-E16 and R-E18)(Fig. 3c). Next, we tested the persistency of the bump during body movement in the darkness. The body movement is simulated by constant activation of the shifter circuits, i.e., steady synaptic inputs to unilateral PEN neurons (see Materials and Methods). During this condition, an accurate HD model should produce smooth and steady and persistent bump movement or a linear trajectory on the bump position vs. time plot. Therefore, we performed linear regression for the trajectory and used the mean *R*^2^ (see Materials and Methods) to measure the motion persistency (Fig. 3d). Our tests indicated that the accuracies of R-E16 and Delta-E16 are comparable, while R-E18 is inferior to the two models, especially when the synaptic weight is strong (Fig. 3e). Delta-E18 was excluded from this analysis because the model could not form a bump in the test condition.

**Fig. 3.**
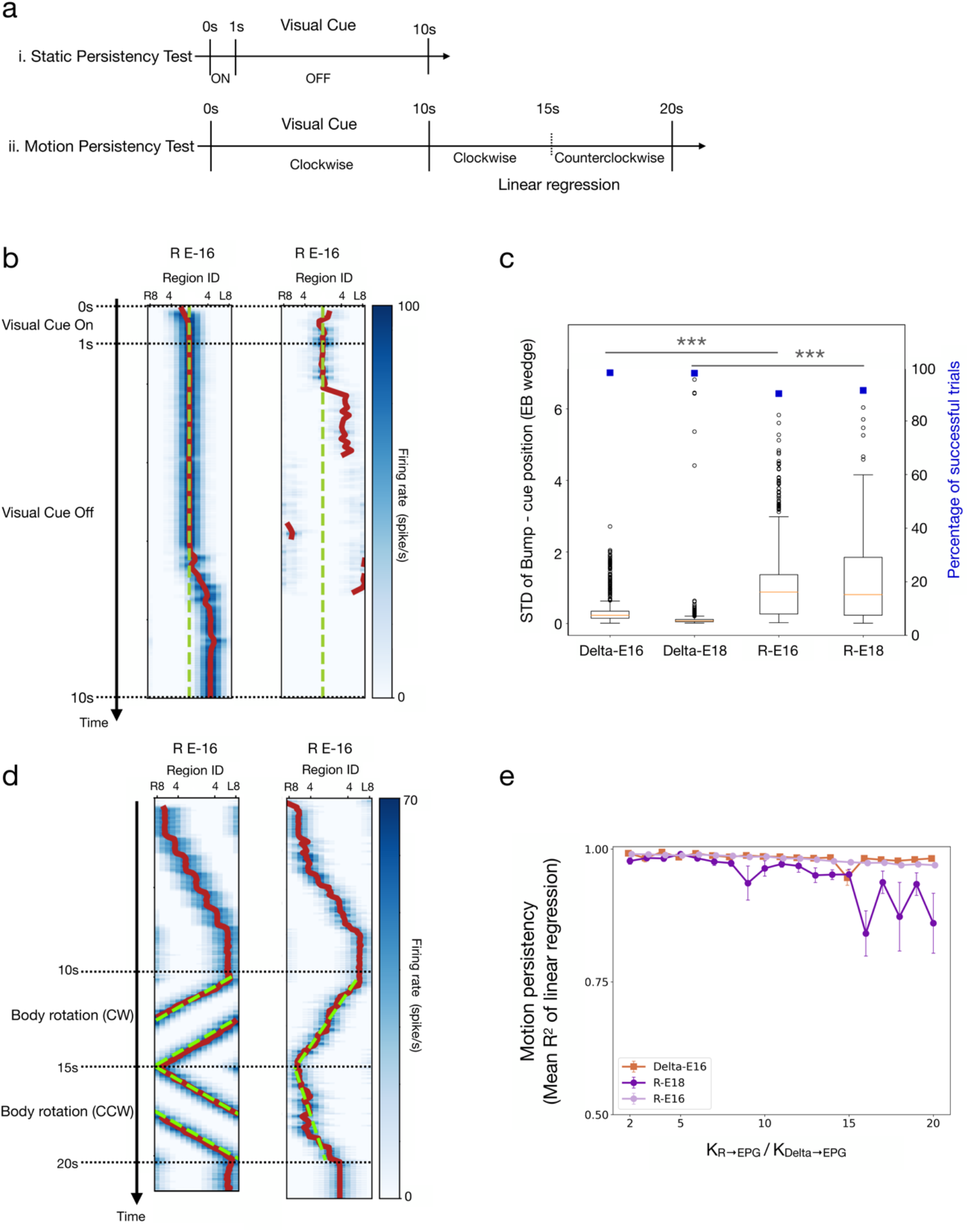
Persistency of the models. **a** The protocols for the static and motion persistency tests (also shown in Fig. 1g). **b** Example trials in the static persistency test. The green dashed lines indicate the visual cue positions, while the red curves represent the peak positions of the bumps. Left: a trail considered successful but with a gradually drifting bump in the last 3s of the trial. Right: a failed trial with a diminished bump. **c** Percentage of successful trials (blue squares) and bump drift (box plots), as measured by the standard deviation (STD) between the bump and cue positions. The mean STDs of the Delta models were significantly smaller than the R models (Wilcoxon rank sum test, ***=p<0.001). **d** Example trials for R-E16 models in the motion persistency test with the linear regression (green dashed lines) of the bump position (red curves). Left: a trial in which the bump shifted smoothly during body rotation, and the trajectory of the bump position could be well fit by a line. Right: a trial in which the bump did not shift smoothly during body rotation, and the linear regression led to a poor fit. **e** Motion persistency, defined as the mean *R*^2^ of the linear regression of the bump trace (see Materials and Methods), as a function of the Δ7/R→EPG synaptic weight for the three models. Delta-E18 was excluded because it could not form a moving bump in the test condition.

### Speed

A good HD model should be able to track the fast azimuthal motion of a visual cue, which drives the bump to shift across EB wedges accordingly. A fast motion causes a brief stimulation period in each region. If the system cannot catch such a fast change, the fast-moving visual stimulus would activate multiple EB regions, and the bump could collapse or spread through the entire EB. Therefore, testing the HD models with different motion speeds of the visual cue is an excellent way to evaluate the capability of the HD models (Fig. 4b). Indeed, our simulations showed that all models could track the visual cue motion at low speeds, but the bump started to spread when the cue speed passed a threshold that depended on the model. We defined the success rate as the percentage of trials in which the bump moved smoothly without spreading and then measured the success rate against the cue speed for all four models. We found that the success rate of each model started dropping at different cue speeds. R-E16 and Delta-E16 could withstand a cue speed up to 0.63*π* rad/s, while the other two models have their success rate drop at lower speeds (Fig. 4c). Therefore, in the speed test, R-E16 and Delta-E16 had comparable performance and were significantly better than the other two models.

**Fig. 4.**
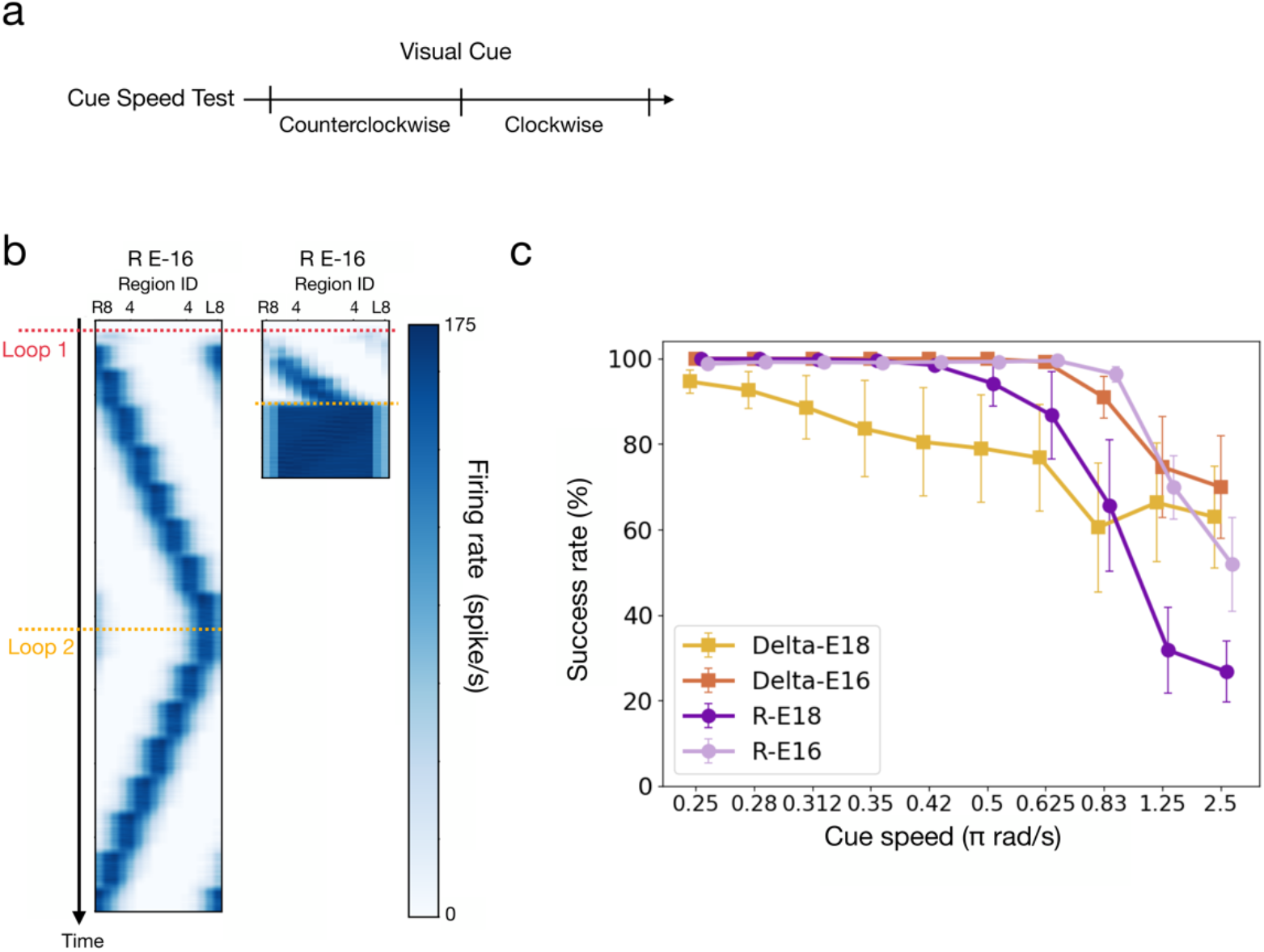
The speed test, which was carried out by examining whether the bump can track a fast-moving visual cue. **a** The protocol for the speed test (also shown in Fig. 1g). **b** Examples of successful and failed trials of the R-E16 model under different speeds of body rotation (left: 0.25*π* rad/s, right: 2.5*π* rad/s) as simulated by shifting the visual cue. **c** The performance of the speed test (measured by the percentage of successful trials) as a function of the movement speed of the cue. The R-E16 and Delta-E16 models had comparable performance and were better than the other two models.

### Dynamical Characteristics

An essential feature of the attractor dynamics of the HD circuits in fruit flies is the ability to move the activity bump accordingly when the visual cue suddenly shifts from one location (azimuthal angle) to another (Kim et al. 2017). Specifically, the bump responds to the cue shift by either “flowing” or “jumping” from the original to the new locations. When the azimuthal distance of the cue shift is small, the bump tends to flow from the original to the new locations. However, the probability of jumping increases with the distance of shift (Kim et al., 2017). In most conditions, we discovered that the bump in all four models could flow from the original to the new cue location, but jumping was much less observed even for a large distance of the cue shift. Therefore, we first focused on investigating the probability of jumping under the condition of a 180° cue shift (Fig. 5b), in which an 80%-100% jumping probability is expected. We randomly selected 750 usable parameter sets from each model and performed 100 trials for each parameter set. A parameter set was regarded as failed if none of the 100 trials exhibited any bump jumping. We found that most parameter sets failed (Fig. 5c). For the parameter sets that did produce jumping, we computed the probability of jumping (out of 100 trials). Delta-E16, R-E16, and R-E18 exhibited comparable jump rates, and the difference is not statistically significant (t-test, p>0.05 for all pairs). Delta-E18 was significantly worse than other models (Fig. 5d). To test how the jump rate change with the shift distance of the visual cue, we selected one representative parameter set from each of the R-E16 and Delta-E16 models, and ran 100 trials for three more distances: 45°, 90°, and 135°. R-E16 exhibited a significantly higher jump rate at the smaller distances (Fig. 5e). At the shift distance of 90°, the jump rates for R-E16 and Delta-E16 were 39% & 15%, respectively. For reference, the observed mean jump rate is ~50% at this distance (Kim et al. 2017).

**Fig. 5.**
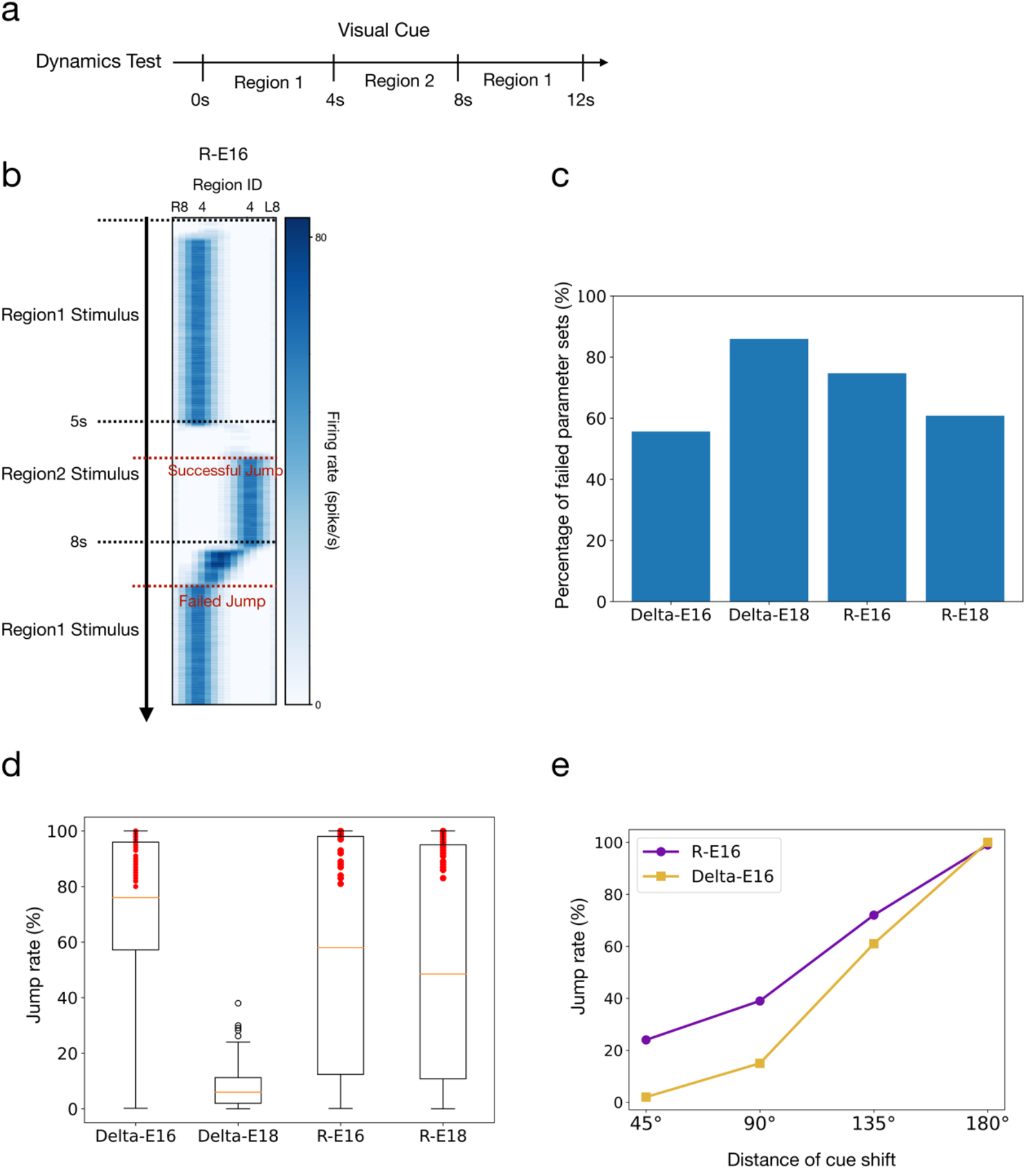
Dynamics characteristics as measured by the probability of bump jumping in response to a sudden shift of the visual cue position. **a** The protocol for the dynamical characteristics test (also shown in Fig. 1g). **b** An example trial showing a successful jump (t=5s) and a failed jump (t=8s) when the visual cue shifted between two EB regions separated by 180°. **c** The percentage of failed parameter sets out of 750 randomly selected parameter sets for each model. A parameter set was considered failed if it did not produce any bump jumping in 100 trials using the protocol as shown in b. The majority of the parameter sets failed in all models. The rest successful parameter sets were selected for further analysis. **d** The jump rates of the successful parameter sets for the four models. The jump rate was defined as the percentage of trials that produced bump jumping. The numbers of parameter sets (red dots) that produce a jump rate higher than 80% are 63, 48, and 33 for Delta-E16, R-E16, and R-E18, respectively. **e** The jump rate as a function of the shift distance of the visual cue for the representative parameter sets from R-E16 and Delta-E16. One hundred trials were performed for each parameter set. The R-E16 performed better than Delta-E16 in this test.

### The Hybrid model

The results presented above show that the R models are more robust, Delta models produce less drift in the static persistency test, while both models perform similarly in other tests. We speculate that the nervous system may implement both mechanisms to combine their advantages and maintain redundancy. We tested this idea by constructing the Hybrid model, which includes both ring neurons and the Δ7 neurons, based on the version of 16 EPG neurons (E16). To determine the best combination of the two inhibitory systems, we swept through different values of *K*_R→EPG_ and *K*_Delta→EPG_ while keeping other parameters as constants based on their best-performed values (*K*_EPG→R_=7, *K*_EPG→Delta_=7, *K*_EPG→PEN_=12.2, *K*_EPN→EPG_=13.6). We performed the Robustness test to select useable parameters and found that the Hybrid model worked with many combinations of *K*_R→EPG_ and *K*_Delta→EPG_ (Fig. 6a). Outside the working regime, the activity bump either spread over all EB, stopped moving, or disappeared. Within the working regime, we selected 11 parameters (red dots in Fig. 6a) that gave rise to more than 80% of the success rates in all of the robustness, speed, and dynamical characteristic tests. We used these parameters in the Hybrid model to compare it with the Delta-E16 and R-E16 models. We first tested how robust the hybrid model is against heterogeneous weight distribution. Following the same procedure described in the robustness test, we added noise to the synaptic weights of all R← →EPG and Δ7← →EPG connections. We found that the Hybrid model was more robust than the R-E16 and Delta-E16 models. The success rate dropped below 0.5 when the STD of the noise exceeded 4.25% of the weights (Fig. 6b). This number is better than the other two models (Delta-E16 = 1.25%, R-E16=1.75%)

**Fig. 6.**
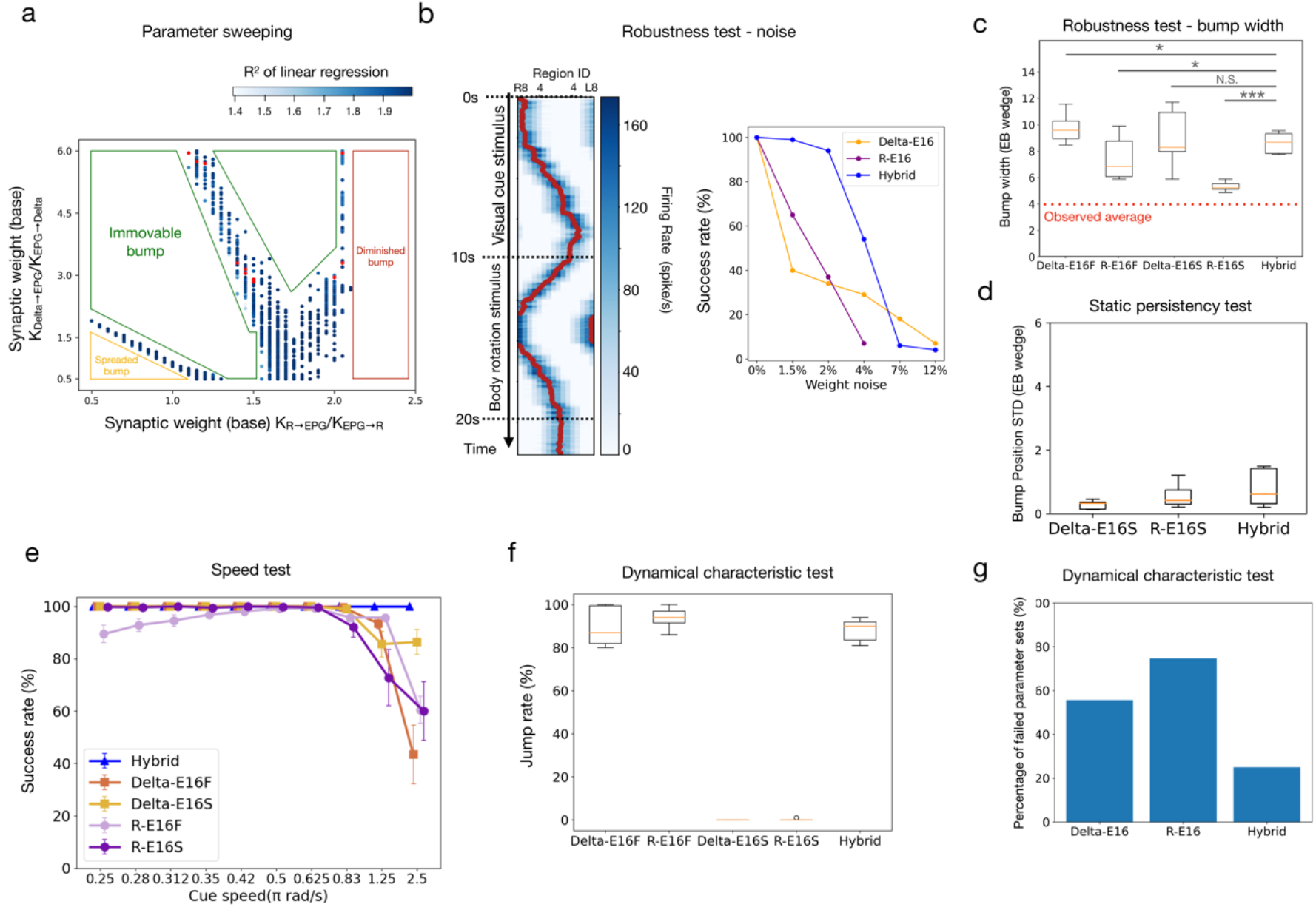
The performance of the Hybrid model, which includes both ring neurons and Δ7 neurons as the inhibitory mechanisms. **a** Usable parameter sets (dots) for the Hybrid model. The saturation of the color indicates the r^2^ score in the motion persistency test. The red dots are the 11 sets of parameters with >80% of success rate in all tests (speed, persistency, and dynamical characteristic) and are selected for comparison with other models in panels c-f. **b** (L) An example activity of the Hybrid model with a noise of 4% of the weights added to all EPG← →R and EPG← →Delta connections. (Right) The success rate of the Robustness test against the noise of different levels. The Hybrid model tolerates more noise than the R-E16 and Delta-E16 models. In all following panels, the Hybrid model is compared with the stable and flexible versions of the R-E16 and Delta-E16 models. A letter (S for stable; F for flexible) is added to the tail of each model name to indicate the version. **c** Bump width. The Hybrid model is not significantly different from the Delta-E16S model (Wilcoxon rank sum test. p-values, Delta-E16F: 0.011, R-E16F: 0.045, Delta-E16S: 0.767, R-E16S: < 0.001) **d** The static persistency test. The Hybrid model is comparable to the stable version of the R and Delta models. The flexible versions of the two models were not shown here because they failed in all trials. **e** The speed test. The Hybrid model outperforms all models at the high cue speed range. **f** The jump rate in the dynamical characteristic test. The Hybrid model performs as good as the Detla-E16F and R-E16F models. The stable versions of the R and Delta models cannot produce any jump. **g** The failure rate in the dynamical characteristic test. The data for delta-E16 and R-E16 models are identical to those shown in Fig. 5c.

Next, we compare the Hybrid model with the Delta-E16 and R-E16 models in the rest tests. However, we discovered that the parameters that led to a better performance in the dynamical characteristics test were generally less stable and performed worse in the persistency test. In comparison, those that led to worse performance in the dynamical characteristics test often performed well in the persistency test.

In fact, the two tests measure the opposite abilities of the activity bump. The persistency test measures the stability of the bump, while the dynamical characteristics test is for the flexibility of the bump. We created a stable version and a flexible version for the R-E16 models (named R-E16S and R-E16F, respectively) by selecting 11 parameters that led to worse and better performance in the dynamical characteristic test, respectively. The same procedure was also performed for the Delta-E16 model (stable version: Delta-E16S, flexible version: Delta-E16F). We found that the bump width of the Hybrid model is comparable to Delta-E16S but worse than the R-E16S (Fig. 6c). For the static persistency test, the performance of the Hybrid model is comparable to the flexible version of the Delta-E16 and R-E18 models, and stable versions failed in all trials (Fig. 6d). In the speed test, the Hybrid model outperformed all versions of the Delta-E16 and R-E18 models (Fig. 6e). In the dynamical characteristic test, the Hybrid model’s performance is comparable to the flexible versions of the Delta-E16 and R-E18 models, while their stable versions failed in this test at all (Fig. 6f & 6g). In summary, except for the bump width, the Hybrid model is comparable to or outperforms the best-performed versions of the Delta-E16 and R-E16 models.

## Discussions

In the present study, we systematically compared five variants (R-E16, R-E18, Delta-E16, Delta-E18, and Hybrid) of the HD circuit models of the fruit flies. The models differ in their global inhibition mechanisms (ring neurons, Δ7 neurons, or both) and the number of EPG neurons (16 vs. 18). Wetested four properties of the models: robustness, persistency, speed, and dynamical characteristics. In general, the models with 16 EPG neurons (R-E16 and Delta-E16) performed much better than those with 18 EPG neurons (R-E18 and Delta-E18). The general performance of R-E16 and Delta-E16 were comparable, but each excelled over the other in different tests. R-E16 performed better in the robustness test, while Delta-E16 was better in the static persistency test. Both performed equally well in the motion persistency, speed, and dynamical characteristic tests. We also noticed that the activity bumps in the Delta models were much broader than the observed bumps, while the widths of the activity bumps in the R models were close to the observed ones. We found that some parameters make R and Delta models more stable (performing better in the persistency test), and others make the R and Delta models more flexible (performing better in the dynamical characteristics test). In other words, the performance in different tests is highly parameter dependent for both models. The Hybrid model combined the advantage of all variances of the R and Delta models and performed very well in all tests with the same set of parameters. Moreover, the Hybrid model can sustain larger heterogeneity in the synaptic weights than the R and Delta models. The only drawback is that it has a bump width similar to that of Delta-E16, which is larger than the observed width.

Why do the E18 models perform worse than the E16 models in most tests? The reason is the disruption of the shifter circuit dynamics caused by the two extra EPG neurons (EPG-R1 & EPG-L1) in the E18 models. The two extra EPG neurons form strong local excitation that induces higher R or Delta7 neuronal activity, which in turn produces stronger inhibition on the PEN neurons. In consequence, the bump gets weakened and even ceases to move.

Several issues need to be considered for the ring and Δ7 neurons as the mechanisms of global inhibition. First, there are many types of ring neurons. Most ring neurons receive inputs from BU (bulb) (Seelig and Jayaraman 2013; Omoto et al. 2017). The rest receives inputs from AOTU (anterior optic tubercle), LAL (lateral accessory lobe), or CRE (crepine) (Hulse et al. 2021). In other words, all ring neurons are projection neurons, not local neurons, as proposed by the classical theories of attractor dynamics. Some of these ring neurons have been shown to deliver spatial information of visual cues to EB (Seelig and Jayaraman 2013; Omoto et al. 2017) or code the wind direction (Okubo et al. 2020). However, the connectomic data (Scheffer et al. 2020) indicate that some ring neurons and EPG neurons form local reciprocal connections, which is exactly what the attractor dynamics needs. Some other studies suggested that when carrying visual information into EB, rather than inhibiting the entire EB, each ring neuron inhibits most EB wedges except for a few (Fisher et al. 2019; Kim et al. 2019). However, based on the connectomic data, many ring neuron classes still form synapses in all EB wedges (Supplemental Fig. 6). Therefore, we suggested that some of the ring neurons exhibit dual functions: a non-uniform inhibition to carry the visual signal into the EPG neurons and a basal signal that provides uniformly global inhibition to EPG neurons. Indeed, a recent study showed that inhibiting or exciting the EPG-innervating R1 neurons led to impaired spatial working memory in the darkness. The result is consistent with the global inhibition hypothesis (Han et al. 2021).

In contrast to the ring neurons, Δ7 neurons are purely local to PB, hence appear to be more suitable to provide global inhibition to the HD circuits. Although each Δ7 neuron only forms patterned inhibition, the entire population of Δ7 neurons provides a global inhibition on the HD circuits. Moreover, blocking neurotransmission in Δ7 neurons reduced the bump’s strength and its head-tracking ability (Turner-Evans et al. 2020). However, the hill-like inhibitory profile formed by individual Δ7 neurons is more complex than what is required by the theories of attractor dynamics, suggesting that Δ7 neurons may also carry other functions. Indeed, a recent study suggested that the Δ7 neurons can assist in creating sinusoidally shaped activity bumps in PFN and PFR neurons, which are part of the circuits that perform vector summation for building an allocentric traveling direction from different sensory cues (Lyu et al. 2021).

As mentioned earlier, ring neurons have been shown to signal visual and other sensory information. Therefore, as a future direction, we will improve the R models by incorporating these functions to examine how non-uniform and uniform inhibition on the EPG neurons may work together to provide multiple functions. The latest connectomic data also reveal that the ring neurons form mutual inhibition (Hulse et al. 2021), a feature that is not included in the present study. This mutual inhibition may make some ring neurons fire stronger than others. The phenomenon may not affect our models because all ring neurons have the same downstream targets. However, when the non-uniform projection of the ring neurons is considered, mutual inhibition will become a critical feature as it could serve as a filtering or selection mechanism for the sensory inputs.

Note that the continuous attractor networks like the ones studied here are generally sensitive to noise or heterogeneity in synaptic weights. However, the large variability in the synaptic numbers within the same type of connections revealed by the connectomic data suggests that the real nervous systems are robust against weight heterogeneity. It has been suggested that short-term synaptic plasticity may play a key role in increasing the stability of the attractor networks against noise and structural heterogeneity (Itskov et al. 2011; Seeholzer et al. 2019). Therefore, we plan to implement short-term plasticity in the future study to increase the robustness of our models.

Finally, the connectomic data reveal abundant dendro-dendritic, axo-axonic, and axo-dendritic synapses in the HD circuits. By reconstructing the 3D structures and implementing a multicompartmental model for each neuron in the circuits, we will be able to accurately simulate the interactions between the neurons and may provide a better insight into the attractor dynamics of the HD circuits.

## Statements and Declarations

The authors declare no conflict of interest.

## Acknowledgment

We thank Dr. Yu-Chi Huang and Mr. Ta-Shun Su for providing the codes used in the present study. This work was supported by National Science and Technology Council grant 111-2311-B-007 −011 - MY3, and by the Brain Research Center under the Higher Education Sprout Project, co-funded by the Ministry of Education and the National Science and Technology Council in Taiwan.

