## Supplemental file for "Global inhibition in head-direction neural circuits: a systematic comparison between connectome-based spiking neural circuit models"

* equal contribution

### Supplemental Fig. 1

**a**

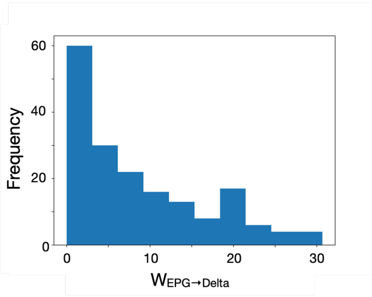

**b**

| 0 | Delta7_L1L9R8_R | Delta7_L8R1R9_L | Delta7_L7R2_L | Delta7_L7R3_L | Delta7_L6R3_L | Delta7_L6R4_L | Delta7_L5R4_L | Delta7_L4R5_R | Delta7_L4R6_R | Delta7_L3R6_R | Delta7_L2R7_R |
|---|---|---|---|---|---|---|---|---|---|---|---|
| EPGt_R9 | 1.33 | 4.00 | 2.00 | 0.00 | 1.00 | 2.67 | 3.13 | 6.30 | 4.25 | 3.25 | 4.78 |
| EPG_R8 | 3.32 | 2.78 | 7.67 | 7.75 | 8.40 | 10.88 | 14.25 | 11.90 | 7.50 | 7.06 | 2.33 |
| EPG_R7 | 3.56 | 13.27 | 9.78 | 14.33 | 10.93 | 19.33 | 11.40 | 5.40 | 2.50 | 2.17 | 3.36 |
| EPG_R6 | 10.00 | 20.33 | 11.78 | 13.67 | 12.27 | 12.00 | 6.79 | 1.50 | 2.00 | 1.67 | 2.62 |
| EPG_R5 | 20.07 | 22.40 | 14.56 | 5.00 | 7.27 | 4.00 | 1.11 | 2.30 | 2.67 | 4.50 | 13.40 |
| EPG_R4 | 21.73 | 14.20 | 4.56 | 1.00 | 1.67 | 3.00 | 2.75 | 3.27 | 16.00 | 12.17 | 21.20 |
| EPG_R3 | 24.60 | 8.50 | 1.57 | 4.50 | 2.43 | 1.50 | 2.86 | 15.87 | 22.67 | 30.67 | 27.33 |
| EPG_R2 | 5.63 | 1.83 | 3.20 | 0.00 | 2.14 | 2.75 | 12.44 | 17.30 | 20.75 | 19.88 | 13.70 |
| EPG_R1 | 1.40 | 4.40 | 2.50 | 4.50 | 6.80 | 22.00 | 27.80 | 26.80 | 28.25 | 24.75 | 10.50 |
| EPG_L1 | 3.75 | 1.25 | 8.50 | 12.50 | 19.90 | 24.25 | 29.00 | 19.90 | 16.00 | 8.88 | 2.00 |
| EPG_L2 | 1.67 | 3.50 | 14.67 | 19.00 | 18.90 | 20.75 | 19.70 | 13.10 | 4.25 | 1.83 | 3.00 |
| EPG_L3 | 6.47 | 17.73 | 20.44 | 19.67 | 20.07 | 18.17 | 9.47 | 2.79 | 3.00 | 2.50 | 2.00 |
| EPG_L4 | 20.07 | 21.00 | 25.89 | 19.00 | 15.67 | 12.50 | 2.58 | 1.91 | 5.20 | 1.50 | 8.00 |
| EPG_L5 | 21.47 | 18.07 | 10.56 | 4.33 | 4.55 | 2.40 | 2.42 | 1.80 | 2.50 | 6.10 | 10.87 |
| EPG_L6 | 13.07 | 7.93 | 2.38 | 2.00 | 2.25 | 1.83 | 3.27 | 2.77 | 7.50 | 8.08 | 6.93 |
| EPG_L7 | 11.00 | 2.54 | 2.88 | 0.00 | 2.22 | 2.00 | 7.64 | 11.00 | 11.17 | 8.17 | 5.87 |
| EPG_L8 | 1.86 | 2.67 | 2.00 | 2.75 | 4.29 | 6.86 | 11.65 | 10.50 | 8.63 | 8.50 | 4.80 |
| EPGt_L9 | 4.33 | 1.33 | 3.25 | 0.00 | 6.25 | 3.00 | 5.00 | 4.40 | 1.00 | 1.00 | 0.00 |

**c**

| Weight threshold | 7 | 8 | 9 | 10 | 11 | 12 | 13 | 14 | 15 |
|---|---|---|---|---|---|---|---|---|---|
| Ng | 91 | 82 | 75 | 71 | 70 | 60 | 52 | 46 | 41 |
| Nc | 26 | 24 | 23 | 23 | 20 | 18 | 17 | 17 | 15 |
| Ng/Nc | 3.53 | 3.41 | 3.26 | 3.08 | 3.5 | 3.3 | 3.05 | 2.7 | 2.9 |

**Supplemental Fig. 1** The connection pattern between EPG and Δ7 neurons and estimation of the dendritic distribution of Δ7 neurons in PB. **a** The distribution of connection weight (the number of synapses) between each EPG (presynaptic) and Δ7 (postsynaptic) neurons. The distribution appears to be bimodal, with a peak at $K_{EPG\to Delta}$=20 and another at $K_{EPG\to Delta} < 3$. **b** The EPG→Δ7 connection table. Each number in a cell represents the mean connection weight between each EPG subtype and Δ7 subtype. There are multiple neurons in each subtype. Each column has 18 cells, which correspond to the 18 glomeruli of PB. We found synapses between every pair of subtypes, but the weight number varies widely. To identify the connection patterns, we set a weight threshold (10 in this case) and only considered the connections with weights above this threshold. After applying the threshold, EPG→Δ7 connections were segregated into clusters (orange boxes). The data are retreived from NeuPrint (https://neuprint.janelia.org/) **c** Based on the distribution shown in **a**, we tested the threshold level between 7 and 15. For each threshold, we counted the number of glomeruli (Ng)(cells in **b**) which exceeded this threshold and the number of clusters (Nc) that contained these threshold-exceeding cells (in **b**). By dividing Ng by Nc, we obtained the average size of a cluster, or the average width of the dendritic distribution of the Δ7 neurons in PB. For the threshold levels we tested, the average size of a cluster fell into the range between 2.7 and 3.5, with a mean of 3.19. Therefore, we used 3 as the size of Δ7 dendritic distribution to construct the Delta-E16 and Delta-E18 models (Fig. 1d).

**Supplemental Fig. 2**

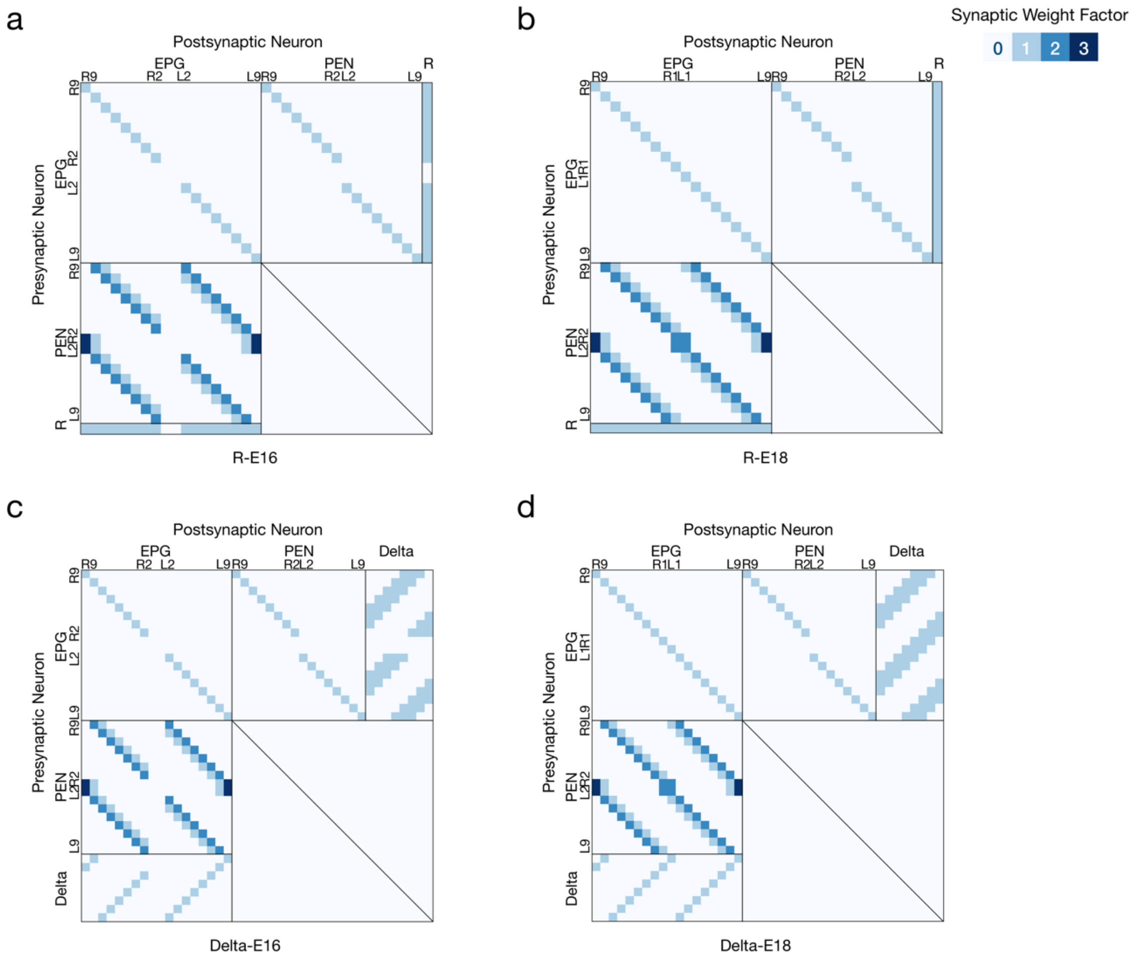

**Supplemental Fig. 2** The connection tables of **a** R-E16, **b** R-E18, **c** Delta-E16 and **d** Delta-E18 models. Note the highly regular and tiling connection patterns in all neuron types. Also, note that EPG_R1 and EPG_L1 only exist in the E18 models but not in the E16 models. The weight factors displayed here have to be multiplied by the tunable weight bases (shown in Supplemental Fig. 5) to become the synaptic weights (in nS).

**Supplemental Fig. 3**

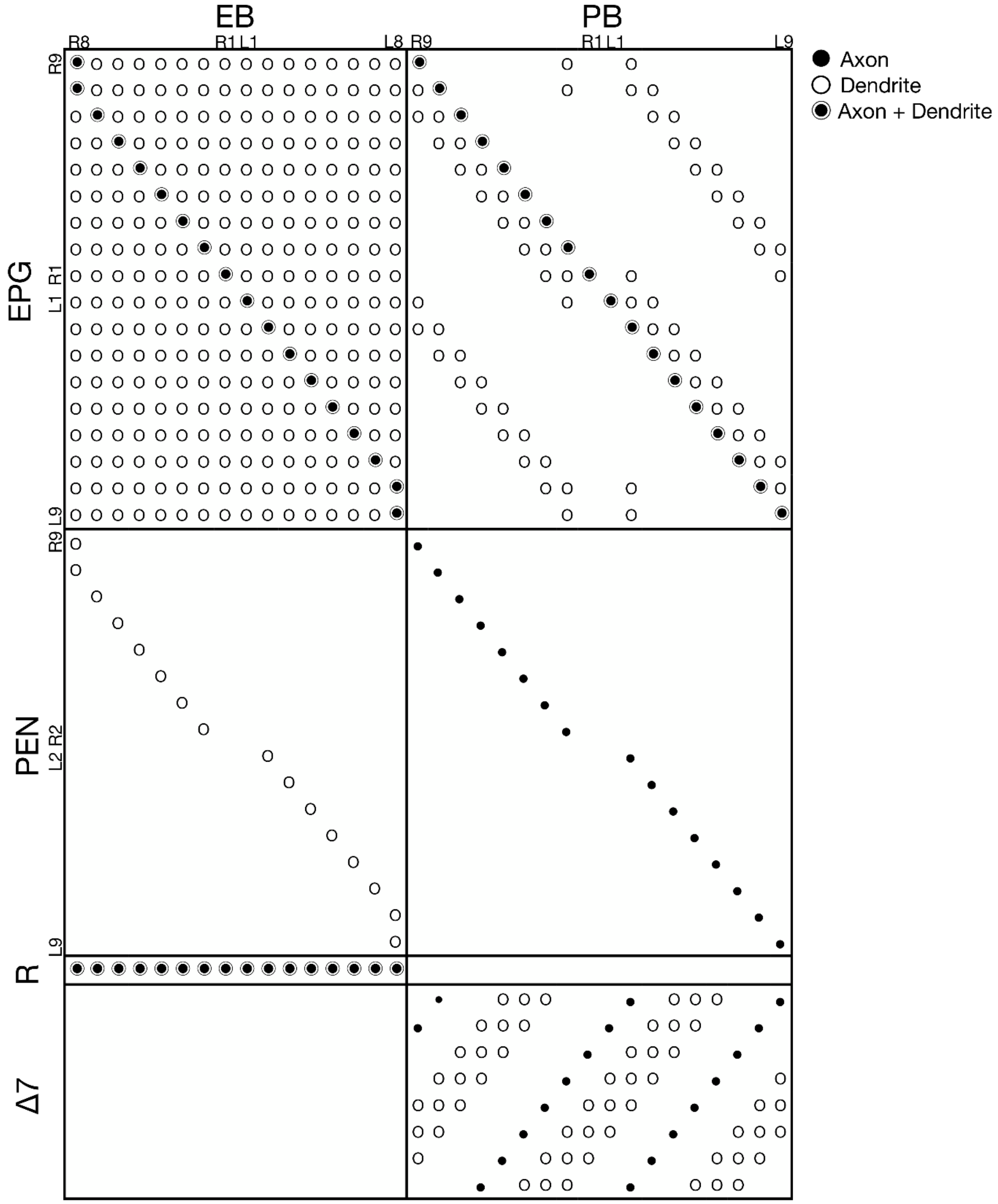

**Supplemental Fig. 3** The innervation tables of the neurons in the models. The Y axis represents neuron subtypes, and the X axis represents the wedges or the glomeruli of EB or PB, respectively.

### Supplemental Fig. 4

a

| | Delta7_L1L9R8_R | Delta7_L8R1R9_L | Delta7_L7R2_L | Delta7_L7R3_L | Delta7_L6R3_L | Delta7_L6R4_L | Delta7_L5R4_L | Delta7_L4R5_R | Delta7_L4R6_R | Delta7_L3R6_R | Delta7_L2R7_R |
|---|---|---|---|---|---|---|---|---|---|---|---|
| EPGt_R9 | 1.33 | 4.00 | 2.00 | 0.00 | 1.00 | 2.67 | 3.13 | 6.30 | 4.25 | 3.25 | 4.78 |
| EPG_R8 | 3.32 | 2.78 | 7.67 | 7.75 | 8.40 | 10.88 | 14.25 | 11.90 | 7.50 | 7.06 | 2.33 |
| EPG_R7 | 3.56 | 13.27 | 9.78 | 14.33 | 10.93 | 19.33 | 11.40 | 5.40 | 2.50 | 2.17 | 3.36 |
| EPG_R6 | 10.00 | 20.33 | 11.78 | 13.67 | 12.27 | 12.00 | 6.79 | 1.50 | 2.00 | 1.67 | 2.62 |
| EPG_R5 | 20.07 | 22.40 | 14.56 | 5.00 | 7.27 | 4.00 | 1.11 | 2.30 | 2.67 | 4.50 | 13.40 |
| EPG_R4 | 21.73 | 14.20 | 4.56 | 1.00 | 1.67 | 3.00 | 2.75 | 3.27 | 16.00 | 12.17 | 21.20 |
| EPG_R3 | 24.60 | 8.50 | 1.57 | 4.50 | 2.43 | 1.50 | 2.86 | 15.87 | 22.67 | 30.67 | 27.33 |
| EPG_R2 | 5.63 | 1.83 | 3.20 | 0.00 | 2.14 | 2.75 | 12.44 | 17.30 | 20.75 | 19.88 | 13.70 |
| EPG_R1 | 1.40 | 4.40 | 2.50 | 4.50 | 6.80 | 22.00 | 27.80 | 26.80 | 28.25 | 24.75 | 10.50 |
| EPG_L1 | 3.75 | 1.25 | 8.50 | 12.50 | 19.90 | 24.25 | 29.00 | 19.90 | 16.00 | 8.88 | 2.00 |
| EPG_L2 | 1.67 | 3.50 | 14.67 | 19.00 | 18.90 | 20.75 | 19.70 | 13.10 | 4.25 | 1.83 | 3.00 |
| EPG_L3 | 6.47 | 17.73 | 20.44 | 19.67 | 20.07 | 18.17 | 9.47 | 2.79 | 3.00 | 2.50 | 2.00 |
| EPG_L4 | 20.07 | 21.00 | 25.89 | 19.00 | 15.67 | 12.50 | 2.58 | 1.91 | 5.20 | 1.50 | 8.00 |
| EPG_L5 | 21.47 | 18.07 | 10.56 | 4.33 | 4.55 | 2.40 | 2.42 | 1.80 | 2.50 | 6.10 | 10.87 |
| EPG_L6 | 13.07 | 7.93 | 2.38 | 2.00 | 2.25 | 1.83 | 3.27 | 2.77 | 7.50 | 8.08 | 6.93 |
| EPG_L7 | 11.00 | 2.54 | 2.88 | 0.00 | 2.22 | 2.00 | 7.64 | 11.00 | 11.17 | 8.17 | 5.87 |
| EPG_L8 | 1.86 | 2.67 | 2.00 | 2.75 | 4.29 | 6.86 | 11.65 | 10.50 | 8.63 | 8.50 | 4.80 |
| EPGt_L9 | 4.33 | 1.33 | 3.25 | 0.00 | 6.25 | 3.00 | 5.00 | 4.40 | 1.00 | 1.00 | 0.00 |

b

| | EPGt_R9 | EPG_R8 | EPG_R7 | EPG_R6 | EPG_R5 | EPG_R4 | EPG_R3 | EPG_R2 | EPG_R1 | EPG_L1 | EPG_L2 | EPG_L3 | EPG_L4 | EPG_L5 | EPG_L6 | EPG_L7 | EPG_L8 | EPGt_L9 |
|---|---|---|---|---|---|---|---|---|---|---|---|---|---|---|---|---|---|---|
| Delta7_L1L9R8_R | 3.20 | 19.20 | 1.83 | 1.00 | 1.90 | 1.93 | 2.07 | 0.00 | 6.57 | 22.50 | 4.88 | 1.00 | 1.69 | 2.22 | 1.60 | 1.00 | 1.67 | 49.10 |
| Delta7_L8R1R9_L | 54.10 | 2.33 | 1.13 | 1.45 | 2.38 | 1.83 | 1.29 | 6.56 | 22.60 | 4.00 | 1.33 | 1.77 | 1.50 | 1.33 | 1.00 | 3.75 | 18.65 | 2.00 |
| Delta7_L7R2_L | 0.00 | 1.00 | 1.17 | 1.00 | 1.83 | 1.00 | 2.75 | 32.17 | 4.00 | 1.00 | 1.60 | 2.56 | 2.50 | 1.00 | 1.00 | 17.67 | 1.50 | 0.00 |
| Delta7_L7R3_L | 0.00 | 1.00 | 1.00 | 1.00 | 0.00 | 0.00 | 23.33 | 2.50 | 1.00 | 2.00 | 4.00 | 2.50 | 2.00 | 1.00 | 4.33 | 13.67 | 0.00 | 0.00 |
| Delta7_L6R3_L | 0.00 | 1.25 | 1.17 | 1.00 | 1.33 | 7.38 | 19.47 | 3.33 | 1.00 | 1.40 | 2.63 | 2.77 | 1.17 | 1.17 | 12.00 | 9.67 | 2.00 | 0.00 |
| Delta7_L6R4_L | 0.00 | 1.00 | 1.00 | 1.00 | 1.00 | 22.67 | 11.33 | 1.00 | 1.50 | 2.50 | 3.00 | 2.50 | 1.00 | 6.00 | 20.67 | 0.00 | 0.00 | 0.00 |
| Delta7_L5R4_L | 0.00 | 1.43 | 1.50 | 1.00 | 3.73 | 11.13 | 1.25 | 1.00 | 1.75 | 2.00 | 2.50 | 2.00 | 4.33 | 19.80 | 6.40 | 1.00 | 0.00 | 1.00 |
| Delta7_L4R5_R | 1.00 | 1.33 | 1.00 | 7.67 | 19.73 | 1.83 | 1.91 | 1.29 | 1.83 | 2.00 | 1.86 | 1.00 | 16.87 | 7.44 | 1.00 | 1.25 | 1.00 | 0.00 |
| Delta7_L4R6_R | 1.00 | 1.00 | 1.00 | 24.17 | 2.25 | 1.75 | 2.00 | 1.00 | 1.67 | 1.50 | 1.00 | 5.75 | 27.50 | 0.00 | 0.00 | 1.00 | 0.00 | 0.00 |
| Delta7_L3R6_R | 0.00 | 1.25 | 4.80 | 16.58 | 1.25 | 2.11 | 4.18 | 1.40 | 2.20 | 1.00 | 9.33 | 19.83 | 3.88 | 1.00 | 1.00 | 1.00 | 1.00 | 0.00 |
| Delta7_L2R7_R | 0.00 | 1.88 | 17.93 | 2.40 | 1.75 | 2.47 | 2.71 | 1.00 | 1.33 | 7.33 | 24.60 | 10.44 | 1.40 | 1.25 | 1.00 | 1.00 | 1.00 | 0.00 |

c

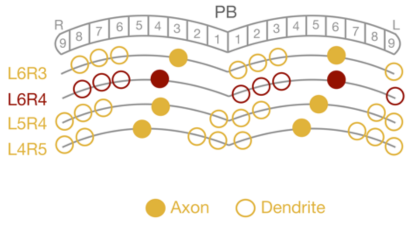

**Supplemental Fig. 4** The connection tables of **a** EPG→Δ7 and **b** Δ7→EPG connections. Data are retrieved from NeuPrint (https://neuprint.janelia.org/). Red rectangles label three "atypical" Δ7 neurons, which were excluded from the presented models. The innervation sites of these neurons are offset from the locations expected for the regular tiling pattern exhibited by other Δ7 neurons. The models did not work if these three atypical neurons were included. **c** A schematic shows how three example "regular" neurons (brown) and one example atypical neuron (purple) innervate the PB glomeruli. The atypical neuron brakes the innervation pattern formed by other regular neurons.

**Supplemental Fig. 5**

a

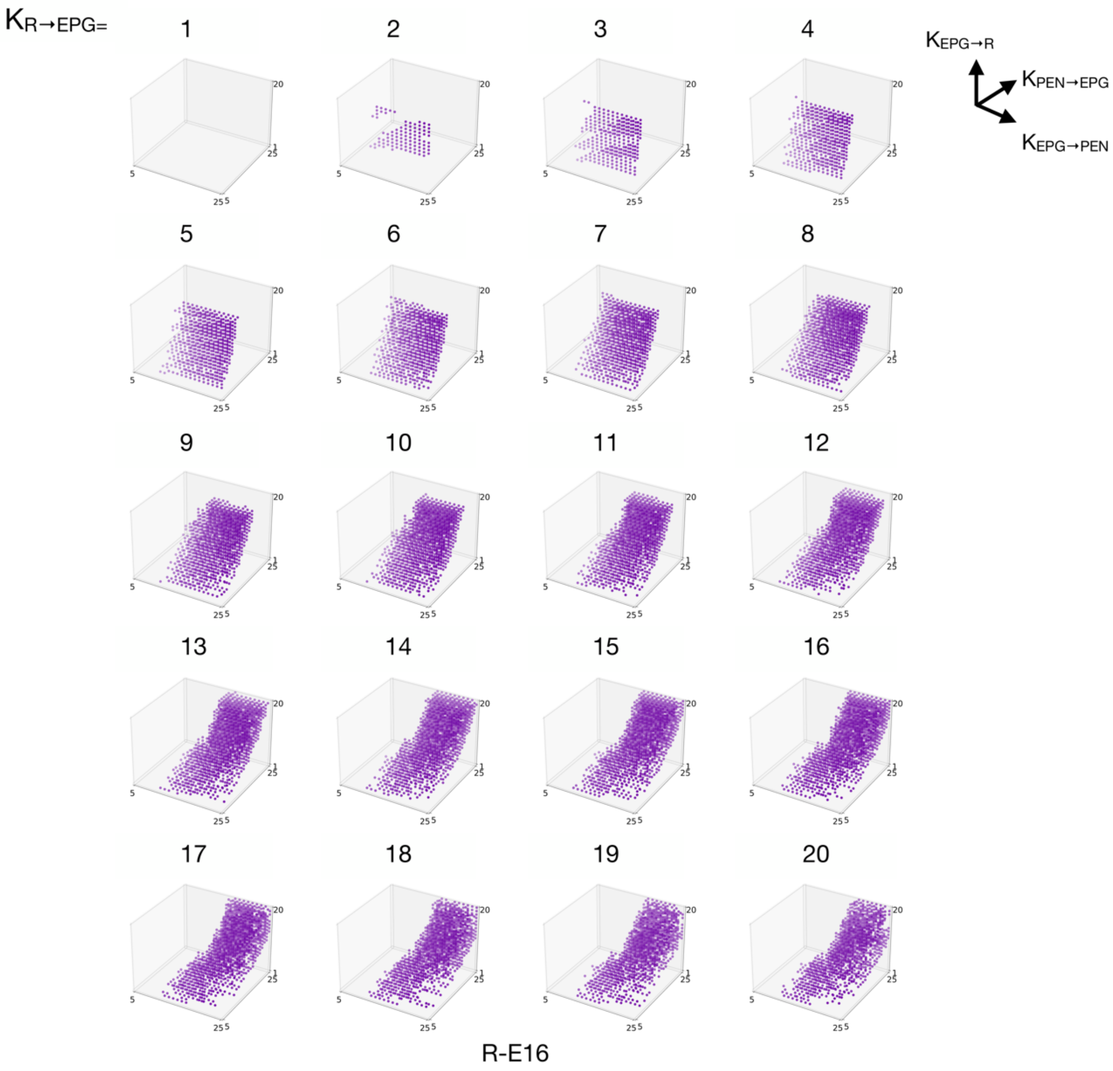

b

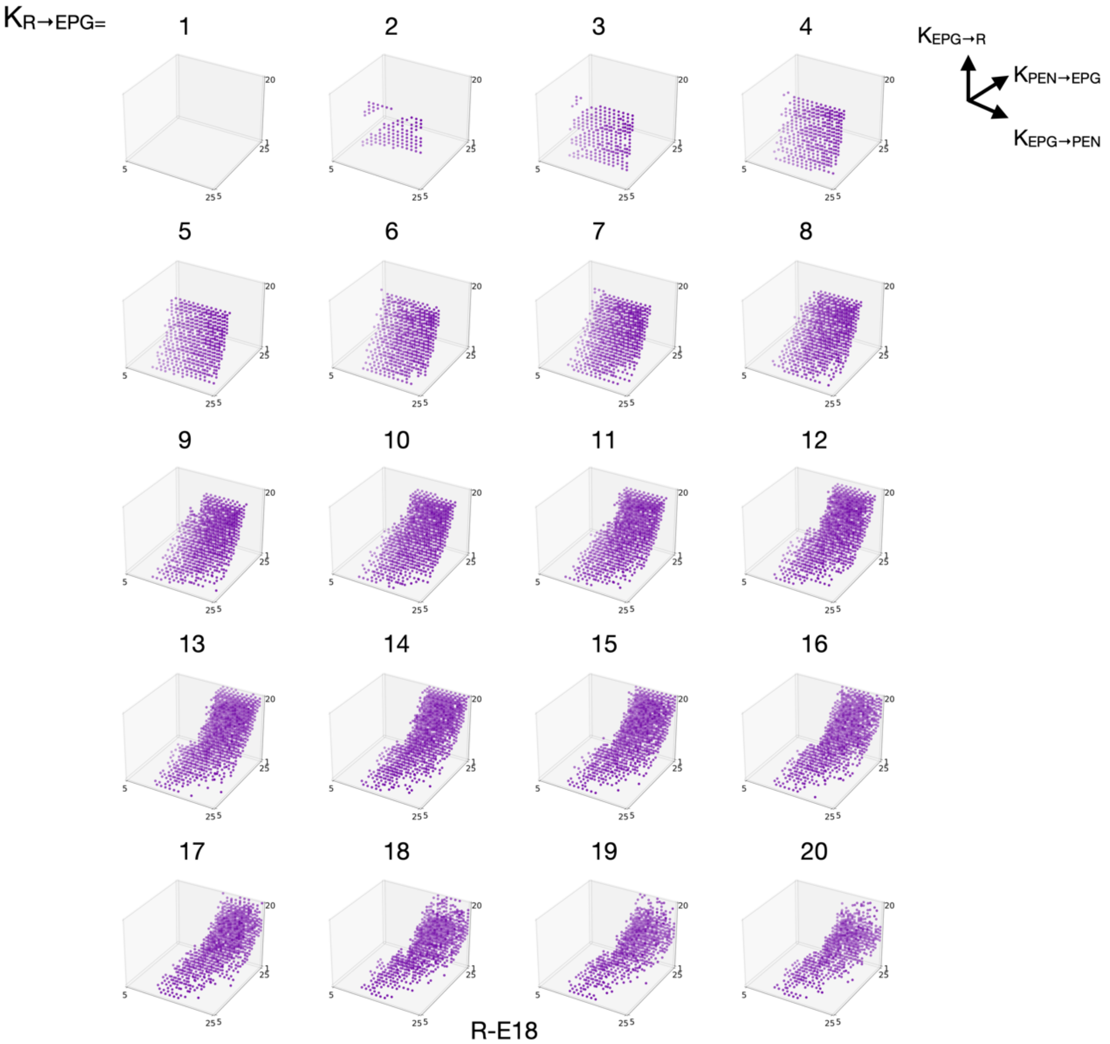

$K_{R \to EPG}$=
$K_{EPG \to R}$
$K_{PEN \to EPG}$
$K_{EPG \to PEN}$

R-E18

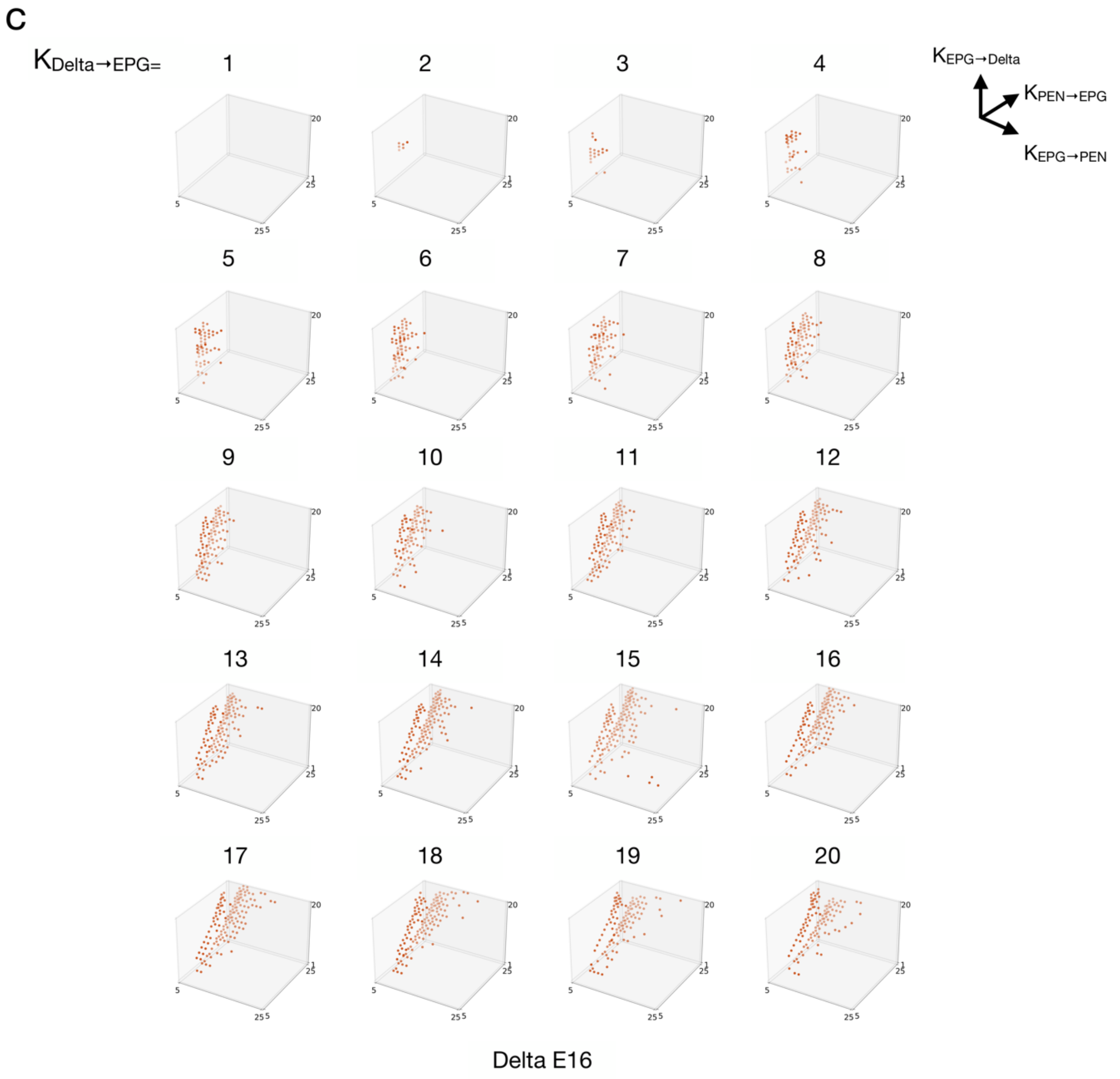

**Supplemental Fig. 5** Parameter sweeping for the weight bases ($K$) in **a** R-E16, **b** R-E18, **c** Delta-E16. Each plot in a panel is a three-dimensional slice for $K_{\mathrm{PEN}\rightarrow\mathrm{EPG}}$, $K_{\mathrm{EPG}\rightarrow\mathrm{PEN}}$, and $K_{\mathrm{EPG}\rightarrow\mathrm{R/Delta}}$ in four-dimensional sweeping. Different plots represent sweeping across $K_{\mathrm{R/Delta}\rightarrow\mathrm{EPG}}$. Each colored dot labels a "usable" parameter set that produced a successful trial in the robustness test. Note that the plots for Delta-E18 are not shown here because no usable parameter could be found for this model in the robustness test.

#### Supplemental Fig. 6

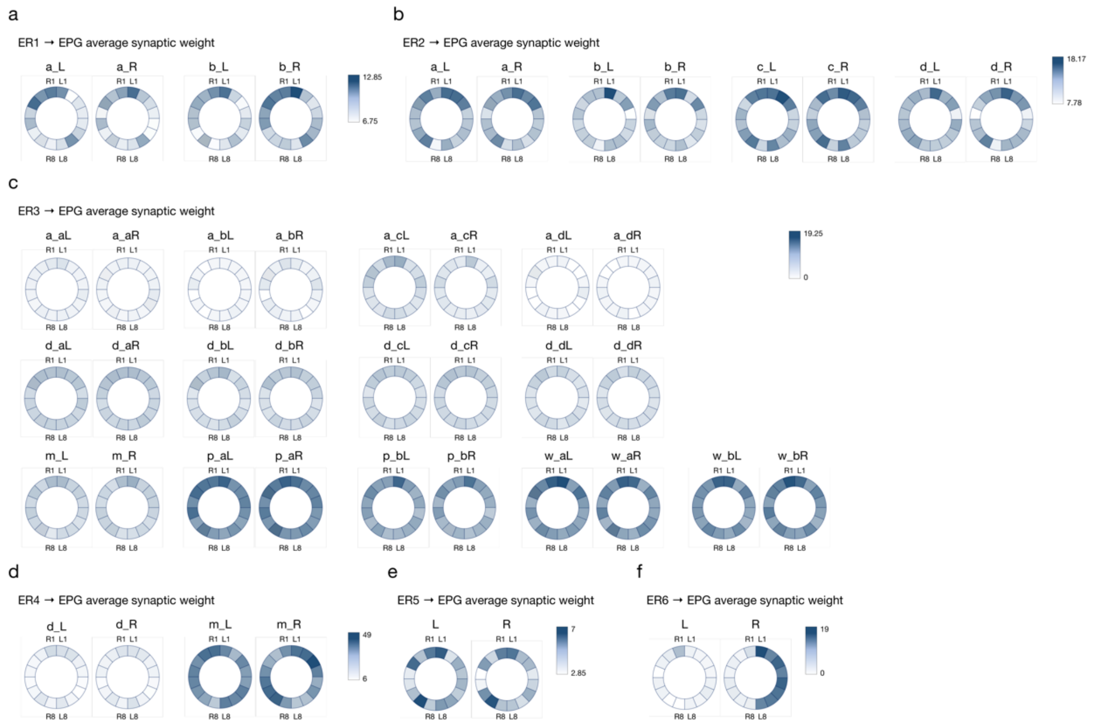

**Supplemental Fig. 6**

The innervation of each type of ring neuron in the EB wedges. The saturation of the color indicates the mean number of synapses a given ring neuron type formed in the given wedge. The data are retrieved from NeuPrint (https://neuprint.janelia.org/).
